# A supervised ontology-aware cell annotation method for single-cell transcriptomic data

**DOI:** 10.64898/2026.01.13.699356

**Authors:** Nimish Magre, Ebtisam Alshehri, Fedor Grab, Yerdos Ordabayev, Steven A. McCarroll, Mehrtash Babadi, Stephen J. Fleming

## Abstract

Many single-cell RNA-seq annotation methods ignore the hierarchical nature of cell type classification. We present a probability propagation strategy that enforces ontological consistency and improves performance when applied to existing models without retraining. Combined with a lightweight logistic regression model trained on 42 million human cells, this yields SOCAM, a fast and interpretable classifier. We also introduce a hop-based F1 score for ontology-aware evaluation. Code and models are available open source.

## Main

Single-cell RNA sequencing (scRNA-seq) enables the measurement of gene expression at single-cell resolution, providing unprecedented insights into cellular heterogeneity that bulk RNA-seq cannot capture [1, 2]. By isolating individual cells, capturing and sequencing their mRNA, and generating a cell-by-gene expression matrix, scRNA-seq has been instrumental in identifying novel cell types, characterizing dynamic states in development and disease, and building reference atlases across tissues and organisms [3, 4]. A key step in scRNA-seq analysis is cell type annotation—the task of assigning biological identities to cells based on their transcriptomes (Fig. 1a).

**Fig. 1.**
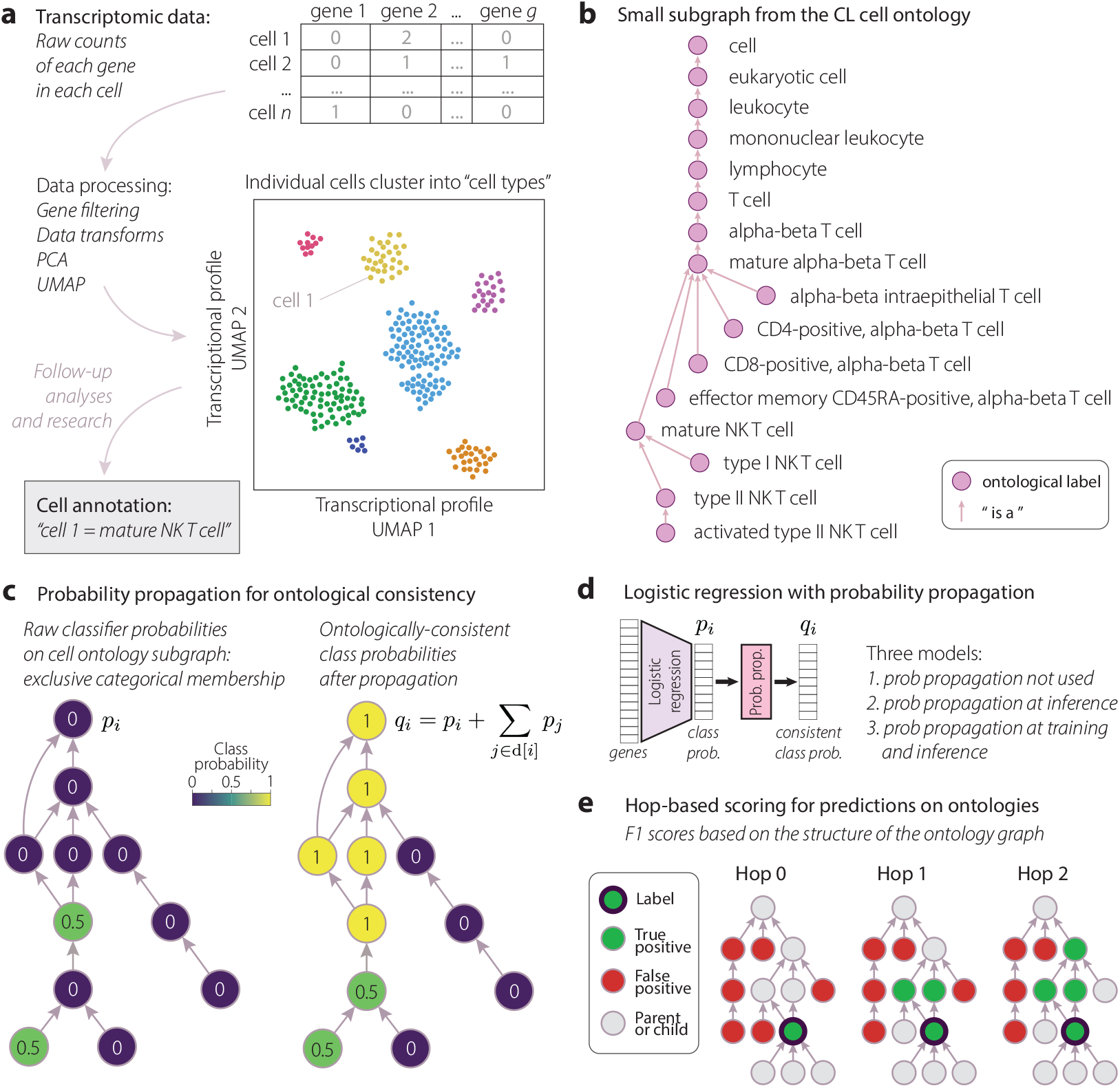
The cell annotation problem and the SOCAM model. (a) Schematic workflow for annotating single cell transcriptomic data with cell type labels. Individual cells cluster based on transcriptional similarity. Clusters correspond to cell types. The object of the “cell annotation” problem is to apply cell type labels to a new dataset. (b) The CL cell ontology organizes cell type labels into a graph. A small subgraph involving “activated type II NK T cell” is shown, leading all the way back to the root node “cell”. (c) Treating the cell annotation task as a naive multi-class classification problem (left) ignores ontological relationships. We introduce a probability propagation strategy to make raw class probabilities consistent with the ontology graph (right): each node contains the sum of itself and all its descendants. (d) The SOCAM model (this study) is a simple logistic regression classifier with probability propagation applied. (e) We introduce an F1 scoring metric based on the ontology graph. Small example graphs are depicted with true positives (green) and false positives (red) for the Hop 0/1/2 F1 scores respectively. The rules for labeling these graphs are described in the Methods section.

Methods such as Azimuth [5], OnClass [6], ScTab [7], and Scimilarity [8] use supervised or semi-supervised learning to classify cells by leveraging labeled reference datasets. While effective, most approaches assume discrete and mutually exclusive cell types and do not fully incorporate the hierarchical relationships encoded in ontologies like the Cell Ontology [9]. In reality, cell types frequently exist on continua defined by overlapping gene expression programs, especially within related lineages or functional modules [10, 11]. For example, studies of hematopoiesis have shown that immune cells transition through gradual transcriptional changes, blurring hard classification boundaries [12]. As a result, assigning a single label to each cell can obscure biologically meaningful relationships and limit the interpretability of annotations.

Supervised classification becomes particularly challenging when categories are structured and overlapping rather than mutually exclusive. The Cell Ontology (CL) provides a standardized, expert-curated framework for cell type definitions and their biological relationships across species [9, 13]. It encodes hierarchical structure through directed edges such as *is_a* (e.g., an “alpha-beta T cell” *is a* “T cell”). Fig. 1b shows a small subgraph from the Cell Ontology, tracing all ancestors of “activated type II NK T cell” and including some additional nodes which are not direct ancestors. (In reality the ontology is not a simple tree structure.) As a result, they may annotate closely related cells using slightly different labels (e.g. “type I NK T cell” versus “mature NK T cell”) without acknowledging how similar or dissimilar these labels are, reducing both robustness and interpretability [9, 14]. This limitation is especially problematic in dynamic or transitional cell states, where ontological context is essential.

To address the challenge of ontologically inconsistent classification, we introduce a lightweight probability propagation strategy that enforces ontological consistency in cell type annotations (Fig. 1c). Starting from raw probabilities output by any classifier, our method distributes probability upward through the Cell Ontology’s *is a* hierarchy, ensuring that parent terms receive probability scores at least as high as their descendants.

Given output multi-class probabilities *p*_*i*_ on the ontology graph nodes *i* ∈ *c*, such that Σ_*i*∈*C*_ *p*_*i*_ = 1, we define the evidence score *q*_*i*_ for node *i* as

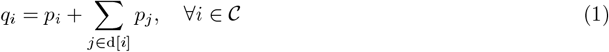

where *c* denotes the set of all cell types in the ontology and d[*i*] denotes the set of all descendants of node The most ancestral root node (“cell” in the Cell Ontology) has *q*_*i*_ = 1. This post-processing step ensures ontological consistency while preserving the probabilistic structure of the output, making it possible to apply this step at inference time without requiring retraining.

To demonstrate the effectiveness of ontology-aware classification, we developed SOCAM (Supervised Ontology-aware Cell Annotation Method), a simple yet performant model built on multi-class logistic regression. SOCAM starts with baseline logistic regression,

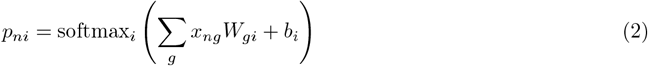

where *p*_*ni*_ is the probability of cell *n* being classified as ontological label *i*, the *g* index denotes genes, *W*_*gi*_ is a learnable weight matrix, *b*_*i*_ is a bias, and *x*_*ng*_ is the measured cell by gene count matrix, appropriately normalized (see Methods).

SOCAM then applies a probability propagation step (Eq. 1) after the softmax, ensuring ontological consistency in predicted cell type probabilities (Fig. 1c). The loss function is:

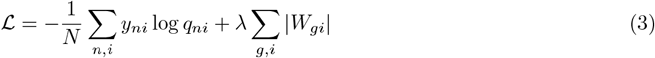

where *y*_*ni*_∈{0, 1} is a one-hot encoding of the true label of each cell *n*, and the term involving *W*_*gi*_ amounts to placing a sparsity-inducing Laplace prior on the weight matrix (controlled by the hyperparamter *λ*).

We remark on three model variants: a baseline logistic regression model without probability propagation, a version with propagation applied only at inference, and a version with propagation during both training and inference. This third variant we refer to as the SOCAM model. Using expression profiles of 20,867 genes as input, SOCAM contains nearly 14 million trainable parameters. We trained the model on 42 million human single-cell transcriptomes from the CZI CELLxGENE Discover platform [15], spanning 670 cell types across 263 tissues and 19 assay types. Unlike conventional classifiers that output flat probability distributions over discrete labels, SOCAM produces probability scores over the Cell Ontology that are ontologically consistent.

To evaluate classification performance in an ontology-aware context, we introduce a **hop-based F1 scoring metric** that accounts for the hierarchical structure of the Cell Ontology (Fig. 1e). Traditional metrics treat only exact label matches as true positives, ignoring the biological relevance of near-miss predictions. Our approach instead defines correctness in terms of ontological proximity: a hop level of 0 considers only an exact match to be a true positive, while hop level 1 expands the true positive set to include the immediate parent(s) of the true label, and so on. This flexible framework enables a graded evaluation of model performance, reflecting the uncertainty and biological continuity often present in fine-grained cell type annotations. By capturing both specificity and ontological relevance, the hop-based F1 score provides a more informative assessment of cell type classification models. Further details on computing hop-based F1 scores can be found in the Methods section.

We evaluated SOCAM and baseline model variants (probability propagation omitted or only applied at inference) on a test set of approximately 600,000 cells from held-out donors, measuring F1 scores across hop levels 0-4 (Fig. 2a). As expected, the base logistic regression model—lacking any ontological context performed worst at all levels. Applying probability propagation at inference improved performance, and the best results were achieved by the full SOCAM model, when propagation was integrated during both training and inference, allowing the model to learn weights that anticipate upward score propagation through the ontology. Further model evaluation stratified by cell type, tissue, assay, suspension type, and sex is presented in Supplementary Fig. 4.

**Fig. 2.**
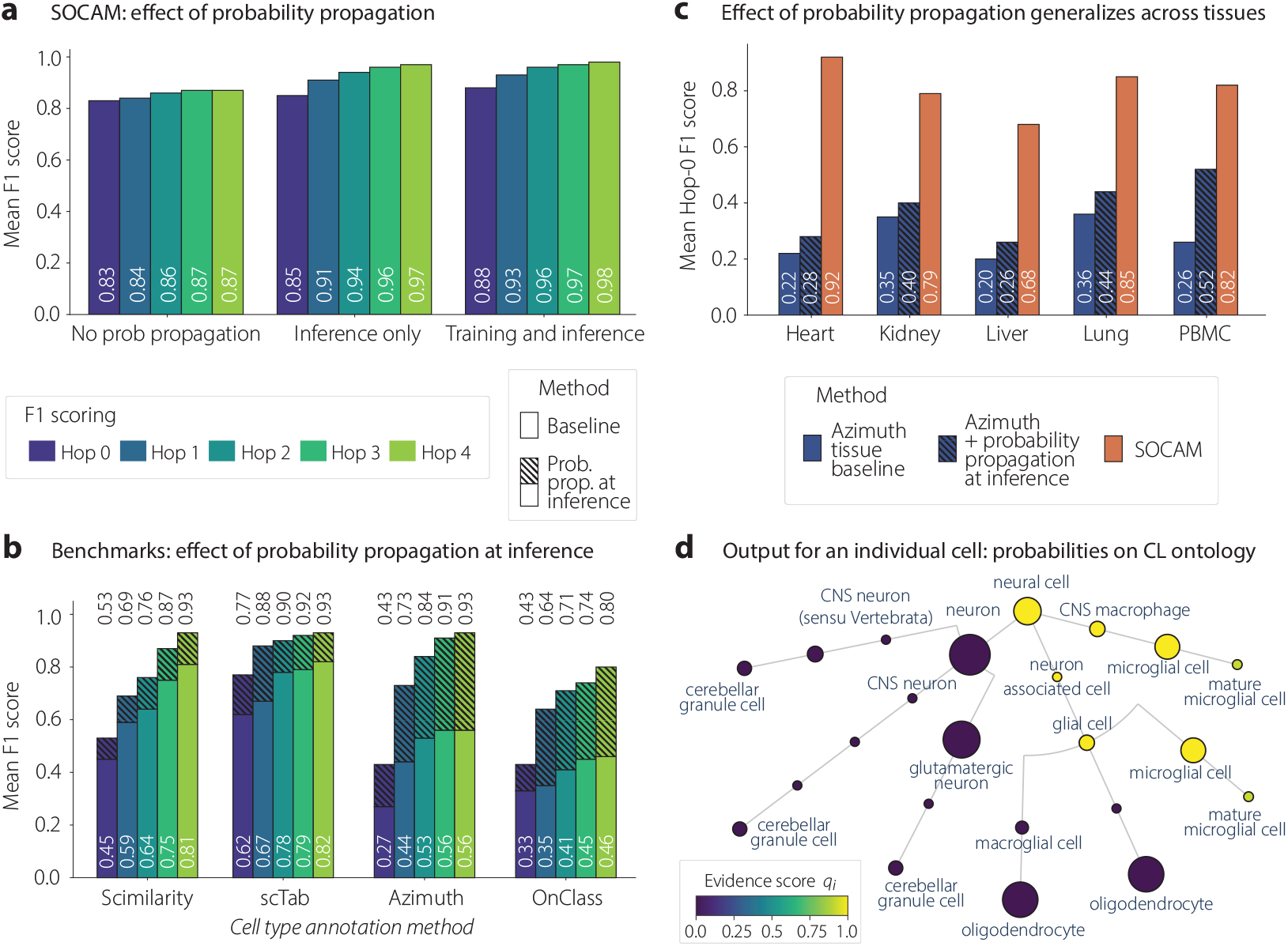
Cell annotation performance by SOCAM and other benchmarks on held-out test data. (a) Mean F1 score at various hop levels for the SOCAM model. Probability propagation was either not applied, applied only at inference time, or applied during training and inference (best performance). (b) Probability propagation can be applied to the outputs of other models at inference time to improve their performance without retraining. Four methods from the literature are benchmarked: Scimilarity, scTab, Azimuth, and OnClass. All methods show marked improvement when probability propagation is applied. OnClass is evaluated on human lung data, as a lung-specific model was the only pretrained model available. All other models are evaluated on all tissues. (c) Azimuth has released tissue-specific annotation models. Probability propagation improves performance for each tissue. SOCAM bar shows performance on the same tissues. (d) Visualization of SOCAM output for a single “microglial cell”. Outputs take the form of consistent evidence scores on the CL ontology graph. We lay a relevant subgraph out here as a tree, so some nodes are repeated. Node size is proportional to the log of the total number of cells with that label in the training data.

**Fig. 3.**
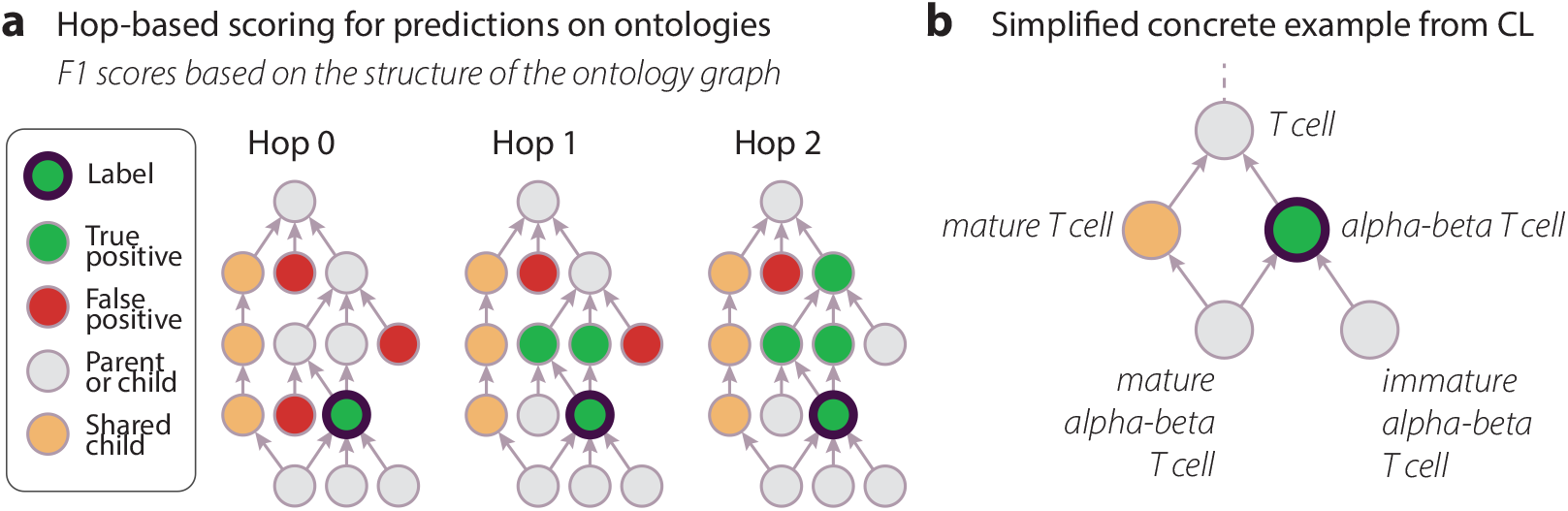
Details of hop-based F1 score calculation. (a) This diagram builds on Fig. 1e and adds the complication of a linkage at the bottom left between a gray child and orange parent. The “shared child” relationship, not present in Fig. 1e, results in excluding the entire orange node branch from being considered false positive. (b) A single concrete example of a “shared child” relationship from the CL ontology. A small subgraph of the CL ontology is shown. From this example it is clear that, given a “true label” of “alpha-beta T cell”, we are not able to call “mature T cell” a false positive label, because a “mature alpha-beta T cell” *is a* “alpha-beta T cell”, and it also *is a* “mature T cell”.

**Fig. 4.**
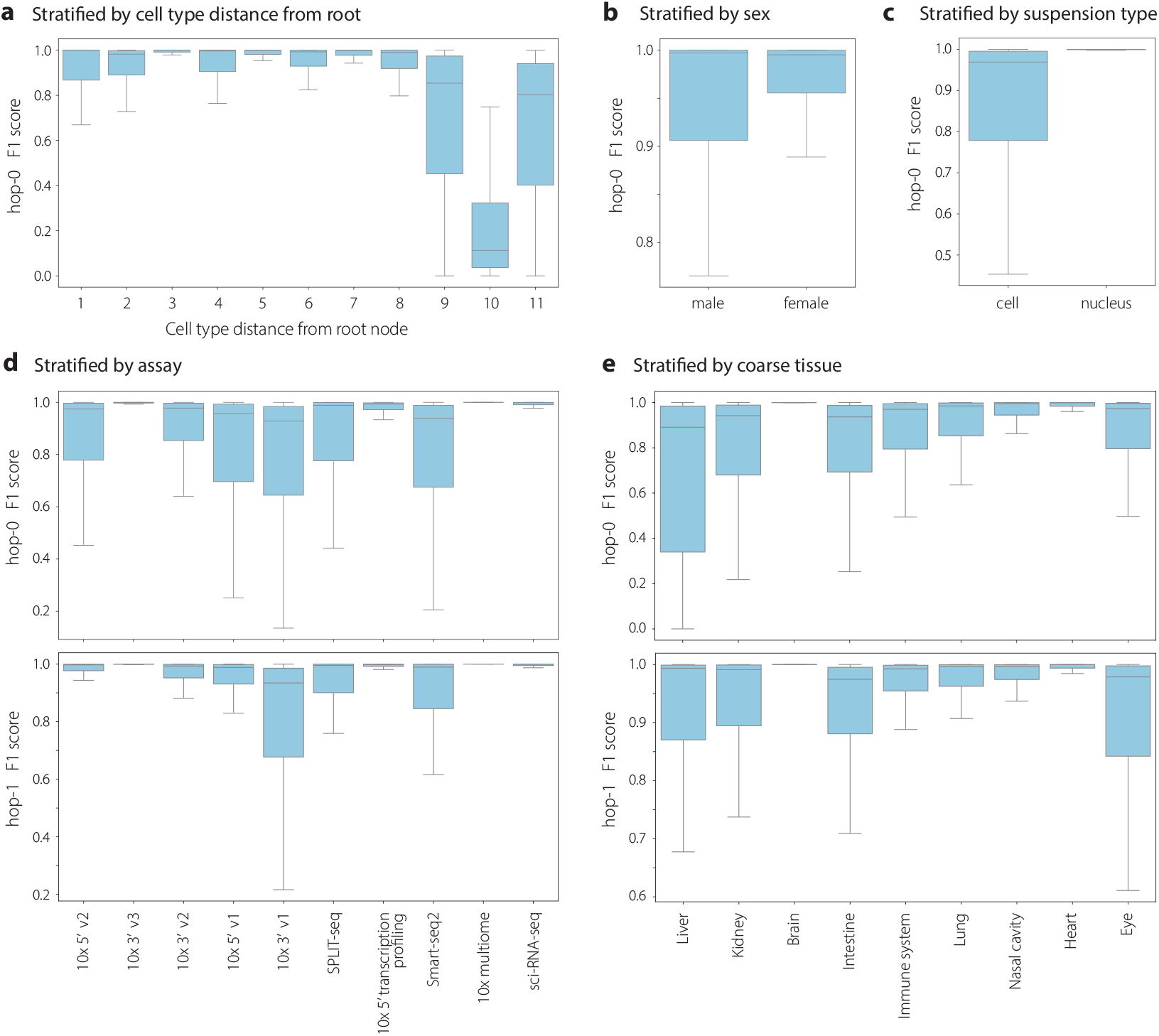
Box plots of hop-*k* F1 score distributions stratified by various validation metadata attributes. (a) Hop-0 F1 scores stratified by target cell type’s distance from the root node (longest distance). Hop scores for coarser cell types tend to have low variance and higher means. (b) Hop-0 F1 scores stratified by the two unique sex metadata attributes. (c) Hop-0 F1 scores stratified by the two unique suspension type metadata attributes. (d) Hop-0 and hop-1 F1 scores stratified by unique assays in the validation set. General trend shows improved mean and variance with increasing hop levels due to propagation. (e) Hop-0 and hop-1 F1 scores stratified by coarse tissue/organ in the validation set. Similar to plots across assays, the general trend shows improved mean and variance with increasing hop levels due to propagation (note y-axis limits).

To demonstrate the generality of this approach, we applied the probability propagation transform to the outputs of four existing annotation tools–Azimuth [5], OnClass [6], scTab [7], and Scimilarity [8]–using pretrained models and a compatible evaluation subset. All methods showed consistent gains in hop-based F1 scores across levels when probability propagation was applied at inference (Fig. 2b), highlighting its value as a plug-in postprocessing step. Notably, tissue-specific Azimuth models showed improved performance across all tissues with propagation (Fig. 2c), and SOCAM achieved strong performance on the same tissue-specific subsets.

By incorporating a lightweight probability propagation step during training and inference, SOCAM outperforms baseline classifiers and the probability propagation step enhances the outputs of existing annotation tools without retraining. The hop-based F1 scoring metric further enables nuanced benchmarking of model performance, particularly in the face of biological ambiguity and annotation uncertainty. SOCAM’s simple logistic regression backbone allows it to scale to tens of millions of cells with modest computational requirements, and its ontology-consistent outputs can be used to help unify annotations across datasets and tissues. We extracted marker gene signatures for all 670 cell types directly from the trained model, clarifying the specific features driving predictions (Methods; Supplementary Table 2).

This study is subject to several limitations. Firstly, the model is trained on a fixed set of gene features and requires test samples to provide counts for the same features. As a supervised linear model trained on human scRNA-seq data, SOCAM’s accuracy at the finest-grained annotation levels is constrained by the availability of high-quality ground truth. Additionally, while probability propagation can be seamlessly applied at inference time to other models, incorporating it during training is less straightforward for nearest-neighbor-based approaches like Scimilarity. Finally, while we benchmarked SOCAM against published models using held-out test data, differences in training data may affect absolute comparisons. Future directions include training on data from more species. The open-source implementation, pretrained model weights, and marker gene lists can be obtained from [https://github.com/cellarium-ai/cellarium-ml/tree/SOCAM].

## Methods

### CL ontology and scRNA reference data

To define ground-truth labels and model predictions for cell-type classification, the systematically structured Cell Ontology (CL) vocabulary is used. The CL nomenclature uniquely identifies each cell type and organizes them according to characteristics, lineage, and functional roles, establishing a hierarchical parent-descendant relationship [9]. Developed under strict quality control and semantic interoperability guidelines by the Open Biological and Biomedical Ontology (OBO) Foundry, the Cell Ontology covers various cell types among different species, providing high-level cell types as mapping points for other species. In addition, it undergoes regular updates to ensure consistency and accuracy. To maintain uniformity across all model variations, the ***2024-01-04*** release was used throughout this project (https://github.com/obophenotype/cell-ontology/releases/download/v2024-01-04/cl.owl).

For reference scRNA data, the ***LTS 2024-07-01*** release from CZ CELLxGENE Discover [15] provides access to over 436 diverse single-cell and spatial transcriptomic datasets, encompassing more than 33 million cells and 2,700+ cell types across human and mouse samples. This extensive dataset supports targeted analyses across various diseases, training sets, and gene expression thresholds, accessible through both web tools and APIs. To train, validate and benchmark all model variations, we make use of the single cell data from human samples representing 670 unique cell types coming from 263 unique tissues, subtissues and 19 assays, ensuring a broad representation of cellular diversity for model training and evaluation. Data from the following assays was excluded for the purposes of this study, due to generally low cell counts: BD Rhapsody Whole Transcriptome Analysis, BD Rhapsody Targeted mRNA, TruDrop, GEXSCOPE technology, STRT-seq, and CEL-seq2. Only cells with “`is primary data`“=True and total UMI counts ≥ 200 were included.

### Gene selection

Genes were initially subset to the 10x GRCh38 reference genes set (https://github.com/cellarium-ai/cellarium-cas/tree/main/cellarium/cas/assets/cellarium_cas_tx_pca_002_grch38_2020_a.json) based on the standard human reference building steps for the 10x Cell Ranger Pipelines (https://www.10xgenomics.com/support/software/cell-ranger/downloads/cr-ref-build-steps#human-ref-2020-a).

The remaining genes were then filtered such that the average per-cell mRNA count per million (often called “TPM”) was less than 1. To implement this filter, we ran the one pass mean var std model from cellarium-ml [https://github.com/cellarium-ai/cellarium-ml] with a total count normalization transform that has a target count of 1 million. After running this model, we filtered out any genes with a mean count value less than 1, resulting in 20,867 genes. A sample config file used to train the one pass mean var std model is available here: [https://github.com/cellarium-ai/cellarium-ml/blob/SOCAM/cellarium/ml/sample_config_files/preprocessing_utility_model_config_files/One_pass_mean_var_std_config.yaml].

### Cell type label filtering

The ontology includes 2914 labels, and we removed all labels which were neither ancestors nor descendants of any cell label found in the training data. This resulted in a total of 2613 target cell types in our simplified cell ontology graph. When training the logistic regression model, we used a weight matrix with label dimension 670, since there are 670 unique cell types for which we have labeled data in the training set. The probability propagation step (Eq. 1) then propagates scores to all 2613 included labels. For label nodes *i* in Eq. 1 that are not included in the regression, *p*_*i*_ is assigned a value of 0. This procedure allows us to compute probability scores *q*_*i*_ on nodes in the ontology graph that were not used by any cell in the training data. During training, loss is computed on the 670 labeled cell type nodes.

### Splitting data into training and evaluation subsets

The CELLxGENE data mentioned above was split into a training set and an evaluation set in the following way. All cells from a subset of individual human donors were chosen to be held out in the evaluation set based on several splitting criteria. We required held-out donors to have (1) the fraction of cell types labeled “unknown” *<* 0.75; (2) average cell type distance from root node ≥ 2; (3) at least 5 unique cell types annotated; (4) at least 500 cells; (5) only one tissue collected; (6) not a rare combination of (tissue, development stage, disease); and (7) a “sex” annotation that is not “unknown”. These filters result in 4789 unique (dataset, donor)s to choose from, each representing a unique individual human donor in CELLxGENE. From this population of donors, we chose 117 in a way that aimed to cover tissues, development stages, sexes, and diseases well. Holding out these 117 donors resulted in holding out about 1.2% of the total cells for the evaluation dataset.

### scRNA-seq data preprocessing

1. Total count normalization: The total UMI count in each cell was rescaled to sum to 10,000 to address variability in sequencing depth. A small epsilon value of 1e-6 was also chosen to avoid any divide by zero errors during the scaling process for this sparse data. The operation is defined by:

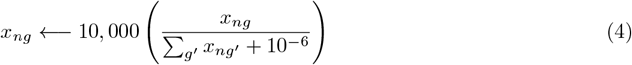
2. Log1p normalization: sc-RNA data often exhibits a heavy-tailed distribution [16] which is mitigated by applying the following transformation:

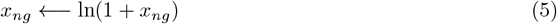
3. Divide by scale: To address the fact that the absolute scale of gene expression is not directly comparable gene-to-gene (e.g. several counts of a transcription factor may signify quite high expression, while several counts of a mitochondrial gene may indicate the gene has unusually low expression), all gene expression values were centered around a common scale using a divide-by-median scaling, where the per-gene median over cells is computed only using nonzero count values *x*_*ng*_:

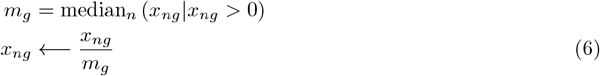

The above median is computed by taking a single pass through all the training data and computing t-digests [17]. We compute these t-digests using the tdigest model available with cellarium-ml [https://github.com/cellarium-ai/cellarium-ml]

### Training details

We train our models using the cellarium-ml framework [https://github.com/cellarium-ai/cellarium-ml]. During training, we use an Adam optimizer [18] with a batch size of 2048 and a learning rate schedule starting at 5 · 10^−7^ with a linear warm-up to 5·10^−3^ for 20% of the total steps, followed by cosine annealing to 0 for the remainder of the steps. We set the hyperparamter *λ* = 100 in Eq. 3 to discourage overfitting. We train each model for 6 epochs (full passes through the dataset). Training was run using Google Cloud Vertex AI pipelines, using a n1-highmem-16 machine and with 2 nvidia-tesla T4 GPUs. Sample config files used to train and test SOCAM are available at [https://github.com/cellarium-ai/cellarium-ml/tree/SOCAM/cellarium/ml/sample_config_files]

### Details on hop scoring

To evaluate the model performance while taking the cell type ontology structure into consideration, we introduce a novel *hop-based F1 scoring metric*. A *hop-level* of 0 defines the “true positive” set as only the target label, while a *hop-level* of 1 additionally includes the set of cell types that are immediate parents of the target cell type, and so on, as shown in Supplementary Figure 3a.

The **true positive rate** for a prediction for cell *n* at hop-*k* is calculated to be:

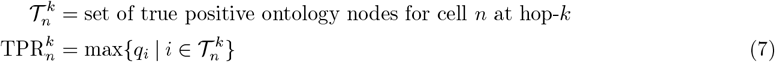

The true positive set 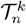 is denoted by the green nodes in Fig. 1e and Supplementary Fig. 3a. At hop-0, the target node alone comprises 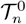, and as the hop level *k* increases, the set of nodes 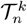 expands to include the direct ancestor nodes of the target node at the corresponding hop level.

Similarly, the **false positive rate** at hop-*k* is calculated as:

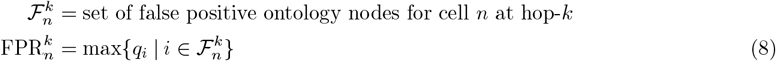

The false positive set 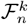 is denoted by the red nodes in Fig. 1e and Supplementary Fig. 3a. We consider a cell type node to be part of the false positive set if (1) it does not have a descendant or ancestor relationship (gray nodes in figure) to the set of cell type nodes that form the true positive set, and (2) it does not share a common descendant with the set of cell type nodes that form the true positive set (orange nodes in Supplementary Fig. 3a).

Supplementary Fig. 3b gives a concrete example explaining the “common descendant” or “shared child” relationship using the CL ontology. Here *alpha-beta T cell* is the target cell type (the author’s label). Direct descendants of the target, *mature alpha-beta T cell* and *immature alpha-beta T cell* in this case, cannot be considered false positives, since a cell labeled *alpha-beta T cell* by an author could be a *mature alpha-beta T cell* or an *immature alpha-beta T cell*. Additionally, if *mature alpha-beta T cell* were an applicable label – a possibility we cannot rule out based on the ontology – then the cell would also be a *mature T cell* (due to the “is a” relationship in the ontology). In general, if a cell type node shares a common descendant with the set of true positive labels, it must be excluded from the false positive set.

Precision and recall at hop-*k* are then computed as:

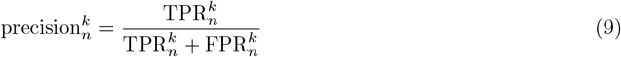

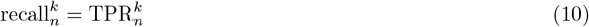

Finally, the F1-score for cell *n* at hop-*k* is computed as:

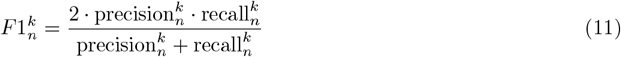

In Fig. 2a-b and Table 1, we report the mean F1 score over cells, mean_*n*_ 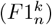, for up to 4 hop levels, since many cell types represented in the validation set are expected to reach the root node (i.e: the “**Cell**” cell type) within 4 hops. Boxplots in the Supplementary Material show distributions of 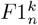 values.

**Table 1.** Performance comparison of probability propagation across methods. Metrics should be compared across methods with caution due to different training data. Metrics uniformly improve when probability propagation is applied at inference (difference between “No propagation” and “Propagation at inference”), without needing to retrain the models. “SOCAM” column refers to SOCAM model outputs benchmarked on the evaluation data subsets used for each of the external methods in the “Method” column.

| Method | Metric | Tissue | No propagation | Propagation at inference | SOCAM |
| --- | --- | --- | --- | --- | --- |
| OnClass | Hop-0 F1 Score | Lung | 0.33 | 0.43 | 0.85 |
|  | Hop-1 F1 Score | Lung | 0.35 | 0.64 | 0.90 |
|  | Hop-2 F1 Score | Lung | 0.41 | 0.71 | 0.93 |
|  | Hop-3 F1 Score | Lung | 0.45 | 0.74 | 0.95 |
|  | Hop-4 F1 Score | Lung | 0.46 | 0.80 | 0.95 |
| scTab | Hop-0 F1 Score | All | 0.62 | 0.77 | 0.88 |
|  | Hop-1 F1 Score | All | 0.67 | 0.88 | 0.94 |
|  | Hop-2 F1 Score | All | 0.78 | 0.90 | 0.96 |
|  | Hop-3 F1 Score | All | 0.79 | 0.92 | 0.97 |
|  | Hop-4 F1 Score | All | 0.82 | 0.93 | 0.98 |
| Azimuth | Hop-0 F1 Score | Heart | 0.22 | 0.28 | 0.92 |
|  | Hop-0 F1 Score | Kidney | 0.35 | 0.40 | 0.79 |
|  | Hop-0 F1 Score | Liver | 0.20 | 0.26 | 0.68 |
|  | Hop-0 F1 Score | Lung | 0.36 | 0.44 | 0.85 |
|  | Hop-0 F1 Score | PBMC | 0.26 | 0.52 | 0.82 |
| Scimilarity | Hop-0 F1 Score | All | 0.45 | 0.53 | 0.88 |
|  | Hop-1 F1 Score | All | 0.59 | 0.69 | 0.93 |
|  | Hop-2 F1 Score | All | 0.64 | 0.76 | 0.96 |
|  | Hop-3 F1 Score | All | 0.75 | 0.87 | 0.97 |
|  | Hop-4 F1 Score | All | 0.81 | 0.93 | 0.98 |

### Obtaining a single best cell type label

The SOCAM model gives probability outputs on the cell ontology for each cell. However, researchers may be interested in obtaining a single, “best” cell type label for each cell. Such a call can be made, however it should be understood that the notion of a “best cell type” call for any given cell is not a well-defined task in general. Coarse-level ontology terms (e.g. T cell, B cell) will have higher relevance scores *q*_*i*_, whereas more granular labels (e.g. CD8-positive, alpha-beta T cell, IgG-negative class switched memory B cell) will have lower scores. The “best cell type” call cannot be equated with the largest score *q*_*i*_, since the node with the largest *q*_*i*_ = 1 is defined to be the root node, “Cell”. In choosing a single “best cell type” label for each cell, there is an inherent trade-off between accuracy and cell type granularity.

Our procedure for choosing a “best cell type” label is to traverse as far through the ontology from the root node as possible while maintaining a relevance score *q*_*i*_ above a specified threshold. After removing cell type nodes with *q*_*i*_ below the provided threshold, we can sort the cell type ontology terms first by distance from root node, and second by *q*_*i*_. We use this sort order to report the top-*k* calls for each cell. Our codebase implements some basic functionality to help users navigate the scored ontology graph and make cell type calls.

### Benchmarking details

We evaluate hop-based F1 scores for four existing annotation tools: Azimuth [5], OnClass [6], ScTab [7], and Scimilarity [8]. While OnClass [6], ScTab [7], and Scimilarity [8] use labels consistent with CL Cell Ontology [9] labels, we filter gene names for each of the pretrained models by first mapping provided gene names to Ensembl gene ids used with SOCAM data [https://github.com/cellarium-ai/cellarium-ml/tree/SOCAM/cellarium/ml/external_benchmarking_details/ensembl_gene_id_to_gene_name_general], then filter SOCAM evaluation data with the genes specific to each of these pretrained models. We then process the SOCAM evaluation dataset with the mapped and filtered genes through each of these pre-trained models to obtain cell type label probability scores.

For Azimuth, we consider only the validation data that has tissue types common to pre-trained Azimuth models. Specifically, we compare outputs across the ***heart, kidney, liver, lung, pbmc*** tissue types. We make use of the ontology references provided by Azimuth [https://azimuth.hubmapconsortium.org/references/] to map prediction outputs to the CL ontology. Missing CL ontology nodes are assigned a *p*_*i*_ = The mappings for tissue-specific labels are available at https://github.com/cellarium-ai/cellarium-ml/tree/SOCAM/cellarium/ml/external_benchmarking_details/Azimuth. We consider probability outputs at all available levels from Azimuth and normalize the results so the total probabilities add up to 1. For cell types available in the CL Ontology but absent from the Azimuth labels, we consider the output probability score to be zero. We provide hop score outputs for Azimuth models with and without the application of probability propagation.

Since Scimilarity performs approximate nearest neighbor search on compact vector representations of scRNA data, the outputs from its nearest neighbor distances are transformed into scores that can then be passed on to the hop based F1 scoring algorithm in the following manner. The distances between a query cell and its *k*-nearest neighbors, *d*_*k*_, are transformed into normalized probability scores through a structured weighting and propagation process. We use 512 neighbors.

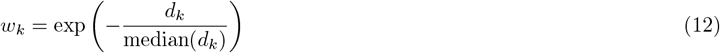

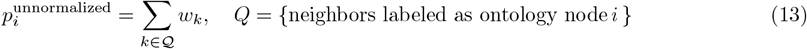

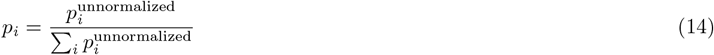

Probability propagation (Eq. 1) is then applied to obtain evidence scores *q*_*i*_ for each ontology node *i*. We further refine the labels by thresholding *q*_*i*_ so that values of *q*_*i*_ *<* 0.1 are set to 0.

### Marker gene identification

We sought to derive biological insights from the trained SOCAM model by identifying the specific genomic features driving its predictions, a process facilitated by the model’s regularized architecture. This analysis serves a dual purpose: it not only elucidates the decision-making process underlying SOCAM’s predictions but also allows us to extract biologically meaningful marker genes for each cell type.

Cells from the combined training and test datasets are aggregated based on cell type and suspension type. This yields a dataset which includes 670 cell types. We perform marker gene analysis on each stratified grouping of cells, taking at most 1000 randomly-chosen cells from each (cell type, suspension type) group. We subsequently identify consensus marker genes by integrating results across suspension types.

To derive interpretable gene-level importance, we utilize integrated gradients [19, 20], a method that attributes the prediction of a model to its input features. For a given cell *n* with a preprocessed (normalized and scaled) gene expression vector *x*_*ng*_, we approximate the integral of the gradients of the model’s output with respect to the input along a linear path from a baseline 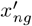 (a zero vector) to the input *x*_*ng*_.

We generate *N* = 100 interpolated inputs 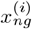 between the zero baseline and the true expression profile:

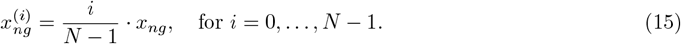

For each interpolated step, we perform a forward pass and computed the gradient of the target propagated cell-type probability, 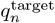, with respect to the input genes using `torch.autograd.grad:`

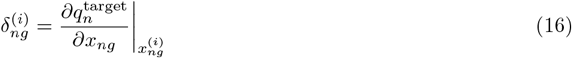

The path integral is approximated by accumulating these gradients, which are then multiplied by the original input expression to derive a raw attribution score, denoted as *a*_*ng*_:

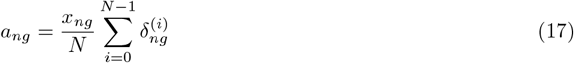

To facilitate comparison across cells, raw attribution scores are normalized to a fixed sum:

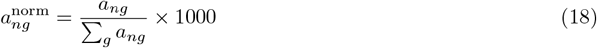

For every cell type, these normalized attribution scores are aggregated across the sampled cells by taking a weighted average based on the target cell type probability predicted by SOCAM, *q*_*n*_. This weighted average down-weights cells which may have been mis-labeled, and which contribute noise to the attribution score estimation.

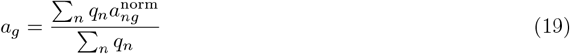

Finally, for a given cell type, genes are ranked by their mean attribution value *a*_*g*_, yielding a data-driven, model-specific marker hierarchy. We provide Supplementary Table 2 which includes a ranking of the top 50 marker genes for each of the 670 cell types. An updated copy of this table is also available in our github repository.

## Data Availability

- [CELLxGENECensus](https://cellxgene.cziscience.com/datasets)
- [List of input genes for the SOCAM model](https://github.com/cellarium-ai/cellarium-cas/blob/main/cellarium/cas/assets/cellarium_cas_tx_pca_002_grch38_2020_a.json)
- [Cell Ontology](https://ontobee.org/ontology/CL)

## Code Availability

Github Repository link: https://github.com/cellarium-ai/cellarium-ml/tree/SOCAM

## Acknowledgements

This work was supported by the Broad Institute’s Stanley Center for Psychiatric Research, the Broad Institute’s Data Sciences Platform, and the National Institute of Mental Health (grants U01MH115727 and UM1MH 130966). The work was also supported by an Escape Velocity grant from the Broad Institute.

## Conflicts of interest

M.B. is a scientific advisory board member at Hepta Bio, and is currently an employee of Isomorphic Labs. Y.O. is currently an employee of Generate:Biomedicines.

## Supplementary Information

### Distribution of F1 scores by cell type distance from the root node

Supplementary Fig. 4a plots the distribution of per-cell hop-0 F1 scores stratified by the ground truth label’s distance from the root node in the cell ontology (longest path) within the validation extracts. These distances encode something about the granularity of a cell type label, since cell ontology nodes closer to the root are, by definition, coarser annotations.

Though the training data comes from multiple donors, assays, and tissues, the overall trend suggests that cell type ontology labels within 8 hops of the root node (86% of nodes seen in the validation set) receive quite high hop-0 F1 scores with tight distributions, emphasizing the effectiveness of the probability propagation process. 76% of cells in the training dataset (92% of cells in validation) were annotated with labels within 8 hops of the root node. It is worth noting that low F1 outliers for cell type distance 1 from root include terms like “abnormal cell”, which is a rare label (37k cells out of 42M) with only 3 descendant nodes, and whose application can be considered somewhat subjective.

As the distance from the root increases, the variance in hop-0 F1 score increases, particularly at 9, 10, and 11 hops from the root node. This trend could be due to the small number of leaf node samples available in the training data as well as heterogeneity in gene expression data for these leaf nodes collected through different assays and representing different tissues.

### Distribution of F1 scores stratified by other metadata

The CZI CELLxGENE database includes additional cell-level metadata such as the assay and tissue. A simple logistic regression model would not be expected to perform equally well across the cell-level metadata attributes, as it should depend on the amount of training data available. In general, the plots show a positive correlation between representation of the metadata characteristic in the training data and the mean hop scores, an inverse correlation between the variance and representation of the metadata characteristic in the training data, and show improved results with increasing hop levels. Supplementary Fig. 4b shows the distributions of hop-*k* F1 scores stratified by sex, while panel (c) shows stratification by suspension type. Supplementary Fig. 4d shows hop-0 and hop-1 F1 scores stratified by assay. The scores improve for all assays as the hop level *k* increases. Supplementary Fig. 4e shows a similar trend when the F1 scores are stratified by tissue.

## Notes

### Competing Interest Statement

M.B. is a SAB member of Hepta Bio, and currently an employee of Isomorphic Labs. Y.O. is currently an employee of Generate:Biomedicines. The remaining authors declare no competing interests.

